# Coevolution of age-structured tolerance and virulence

**DOI:** 10.1101/2023.12.18.572099

**Authors:** Lydia J. Buckingham, Ben Ashby

## Abstract

Hosts can evolve a variety of defences against parasitism, including resistance (which prevents or reduces the spread of infection) and tolerance (which protects against virulence). Some organisms have evolved different levels of tolerance at different life-stages, which is likely to be the result of coevolution with pathogens, and yet it is currently unclear how coevolution drives patterns of age-specific tolerance. Here, we use a model of tolerance-virulence coevolution to investigate how age structure influences coevolutionary dynamics. Specifically, we explore how coevolution unfolds when tolerance and virulence (disease-induced mortality) are age-specific compared to when these traits are uniform across the host lifespan. We find that coevolutionary cycling is relatively common when host tolerance is age-specific, but cycling does not occur when tolerance is the same across all ages. We also find that age-structured tolerance can lead to selection for higher virulence in shorter-lived than in longer-lived hosts, whereas non-age-structured tolerance always leads virulence to increase with host lifespan. Our findings therefore suggest that age structure can have substantial qualitative impacts on host-pathogen coevolution.

## 1. Introduction

Pathogens have evolved a wide range of strategies to spread within and transmit between their hosts, often causing significant damage to their hosts in the form of mortality or sterility virulence. Hosts have, in turn, evolved defences against pathogens. Defences may be in the form of resistance, which has an adverse effect on the pathogen (preventing infection and/or subsequent reproduction and transmission of the pathogen) or in the form of tolerance, which does not reduce pathogen fitness (mitigating the negative consequences of pathogen reproduction and transmission for the host). Empirical evidence suggests that tolerance is an important mechanism in protecting a variety of hosts, including plants (Pagan and Garcia-Arenal 2020) and animals (Raberg et al. 2007), from the adverse effects of pathogens. The coevolution of host tolerance and pathogen virulence is therefore likely to have important implications for both epidemiology and evolutionary biology (Little et al. 2010; Seal et al. 2021).

Empirical evidence suggests that host responses to infectious disease vary with host age (Jarosz and Burdon 1990; Sait et al. 1994; Glynn and Moss 2020). However, to measure the level of susceptibility of a host to pathogens, empirical studies often only consider disease-induced mortality or the severity of symptoms of the disease, and so this measure of susceptibility is likely to include components of both resistance and tolerance. However, some empirical work has sought to isolate the effects of resistance and tolerance mechanisms and subsequently looked specifically at age-specific tolerance (Ramsden et al. 2008; Jackson et al. 2014; Regoes et al. 2014; Sorci et al. 2021). Variation in tolerance has been observed between adults and older individuals (Regoes et al. 2014; Sorci et al. 2021) but also between juveniles and adults (Ramsden et al. 2008; Jackson et al. 2014).

There are a number of reasons why hosts may experience changes in tolerance as they age. One possible explanation is that the pathogen may inflict different levels of virulence on hosts of different life-stages. This might generate greater selection for tolerance in the life-stage during which the pathogen is more harmful, which could in turn lead to the evolution of age-structured tolerance. Alternatively, hosts may be exposed to more pathogens as juveniles than as adults, or vice versa, perhaps due to differences in social behaviour, which may also lead to variation in selection on tolerance at different life-stages. Prior exposure and immune priming may also be factors; older hosts are more likely to have been exposed to the pathogen previously and so may have greater tolerance than younger hosts. Another possibility is that the host pays different costs for exhibiting tolerance at different life-stages, perhaps due to resource allocation constraints. For example, it may be advantageous to invest in tolerance as a juvenile but not as an adult if that would cause resources to be diverted away from reproduction.

Multiple studies have identified age-specific trade-offs between disease susceptibility and reproduction (Chaplin and Mann 1978; Simons 1979; Tian et al. 2003). For example, Tian *et al*. (2003) found that a gene which reduces susceptibility to a bacterial pathogen in *Arabidopsis* is also associated with a reduction in seed set (Tian et al. 2003). However, as susceptibility to the disease is measured by observing symptoms in hosts exposed to the pathogen, this may be indicative of either resistance or tolerance, or a combination of both. Elsewhere, host tolerance specifically has been found to be affected by diet, which suggests that resource allocation may impact on tolerance levels (Budischak and Cressler 2018; Garland et al. 2022). Tolerance to one pathogen has also been found to trade off with tolerance to another (Montes et al. 2020). Therefore, increasing tolerance to one pathogen may impact on reproduction or survival (depending on whether other pathogens in the environment inflict mortality or sterility virulence).

There is also empirical evidence to suggest that immune responses to pathogens can evolve independently at different life-stages (Bruns et al. 2022), although juvenile and adult tolerance traits may be strongly correlated due to shared immune mechanisms. Therefore, juvenile and adult tolerance may be independent traits under contrasting selection pressures, or tolerance may be a lifelong trait with no differentiation between the juvenile and adult stages (or juvenile and adult tolerance may be partially correlated). Theoretical models of tolerance evolution generally assume that tolerance is a single, lifelong trait (for example (Boots and Bowers 1999; Roy and Kirchner 2000; Restif and Koella 2004; Miller et al. 2005; Best et al. 2014)).

In real host-pathogen systems, the pathogen would generally be expected to evolve at least as quickly as the host. It is therefore important to consider coevolution between the host and pathogen, as opposed to evolution only in the host. Coevolution between host tolerance and pathogen virulence has been studied previously in non-age-structured populations (Best et al. 2008, 2010, 2014), including by Best *et al*. (2014), who found that these models do not typically generate diversity through polymorphism or cycling (Best et al. 2014). The evolution of host resistance in age-structured populations has been theoretically explored (Ashby and Bruns 2018; Buckingham et al. 2023; Buckingham and Ashby 2023), but as far as we are aware, the (co)evolution of age-structured tolerance and virulence has yet to be studied.

In this paper, we theoretically investigate how age-structure affects the coevolution of host tolerance and pathogen mortality virulence. Specifically, we consider the effects of juvenile versus lifelong tolerance on coevolutionary dynamics. We find that coevolutionary cycling only occurs when tolerance is restricted to the juvenile stage and that age-structure can impact upon the qualitative effects of varying lifespan, causing pathogen virulence to be higher in shorter-lived than in longer-lived hosts.

## 2. Methods

### 2.1 Model description

We consider a model for the coevolution of host tolerance and pathogen mortality virulence in an asexual, well-mixed host population structured by age into juvenile (*J*) and adult (*A*) stages. Let *S*_*i*_ and *I*_*i*_ be the densities of susceptible and infected hosts respectively at life-stage *i* ∈ {*J, A*}, giving a total host population density of *N* = *S*_*J*_ + *S*_*A*_ + *I*_*J*_ + *I*_*A*_. Juveniles mature into adults at rate *g* > 0 and adults reproduce at a maximum rate *a* > 0 subject to density-dependent competition given by *q* > 0 (juveniles do not reproduce). Juvenile and adult hosts die naturally at rates *b*_*J*_ and *b*_*A*_. Disease transmission is assumed to be density-dependent, with transmission rate *β* and force of infection (rate at which susceptible hosts become infected) given by *λ* = *β*(*I*_*J*_ + *I*_*A*_). Hosts recover from infection at a constant rate *γ*. Mortality virulence is given by *α* > 0 (the disease-associated mortality rate) and is reduced due to tolerance by a factor of (1 − *τ*_*J*_) and (1 − *τ*_*A*_) in juveniles and adults respectively. We seek to determine the effect of age-structured tolerance on the coevolution of tolerance and virulence and so we consider separately the cases where juvenile tolerance evolves with adult tolerance fixed and where juvenile and adult tolerance are equal.

In a monomorphic population and in the absence of any costs of tolerance or virulence, the population dynamics are described by the following set of ordinary differential equations (see Fig. 1A for model schematic and Table 1 for a list of model parameters and variables):

**Table 1:** Model parameters and variables.

| Parameter/<br>variable | Description | Default value or<br>range |
| --- | --- | --- |
| $a$ | Reproduction rate of adult hosts | n/a |
| $a_0$ | Baseline reproduction rate of adult hosts | 5 |
| $b_J, b_A$ | Natural mortality rate of juvenile/adult hosts | 1/5 |
| $b$ | Natural mortality rate of all hosts when $b_J = b_A$ | 1/5 |
| $c_1$ | Strength of tolerance-reproduction trade-off | 0.25 |
| $c_2$ | Shape of tolerance-reproduction trade-off | 3 |
| $g$ | Host maturation rate | 1 |
| $I_J, I_A$ | Density of infected juveniles/adults | n/a |
| $N$ | Host population density | n/a |
| $q$ | Strength of host density-dependence | 1 |
| $S_J, S_A$ | Density of susceptible juveniles/adults | n/a |
| $t$ | Time, measured in arbitrary units | n/a |
| $\tau_A$ | Adult tolerance | 0 |
| $\tau_J, \tau_L$ | Juvenile/lifelong tolerance | $0 \leq \tau_J, \tau_L \leq 1$ |
| $\alpha$ | Mortality virulence | $0 \leq \alpha$ |
| $\beta_0$ | Baseline transmission rate | 10 |
| $\gamma$ | Recovery rate | 0 or 1 |
| $\lambda$ | Force of infection | n/a |
| $\phi$ | Relative pathogen mutation rate | 20 |

**Fig. 1.**
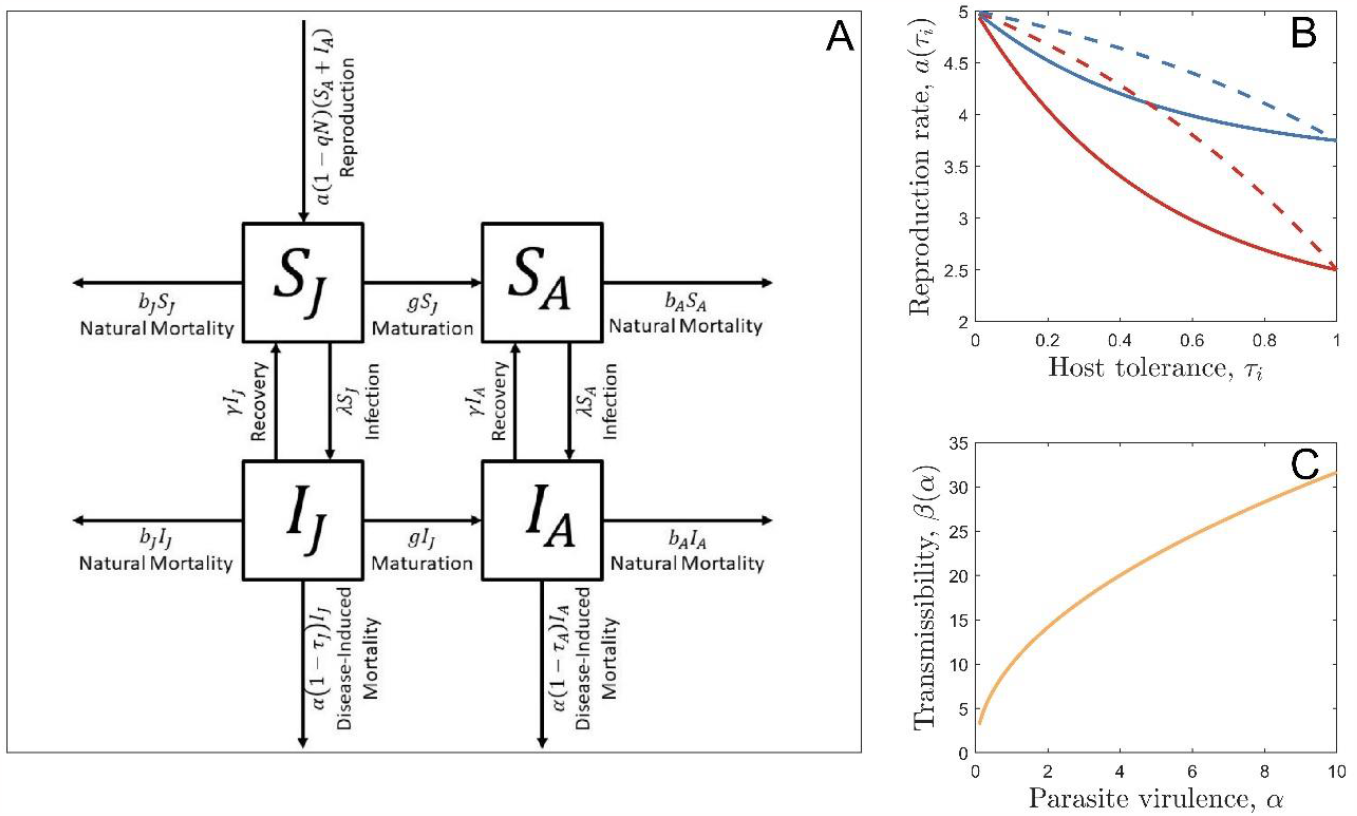
(A) Model schematic for a monomorphic population. (B) Examples of tolerance-reproduction trade-offs (with *a*_0_ = 5). Trade-off strength is controlled by the parameter *c*_1_; a relatively strong trade-off (*c*_1_ = 0.5, red) results in a much larger reduction in the birth rate for a given level of tolerance than a relatively weak trade-off does (*c*_1_ = 0.25, blue). Trade-off shape is controlled by the parameter *c*_2_; a positive value (*c*_2_ = 2, solid) means that the costs decelerate as tolerance increases whereas a negative value (*c*_2_ = −1, dashed) leads to accelerating costs. (C) The virulence-transmission trade-off, with *β*_0_ = 10.

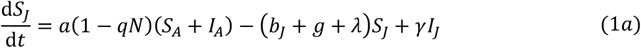

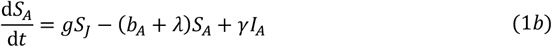

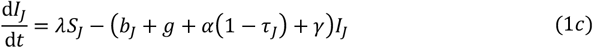

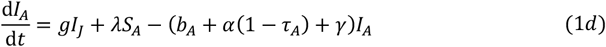

We consider two model versions, one in which tolerance is a lifelong host trait (*τ*_*J*_ = *τ*_*A*_ = *τ*_*L*_) and another in which juvenile tolerance can evolve and adult tolerance is fixed. It is possible to non-dimensionalise this model by rescaling population sizes by 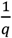 and time by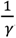; we can therefore fix *q* = 1 and *γ* = 1 in equation (1) without loss of generality (see *Online Resource*). However, we also consider the case where there is no recovery from infection, in which case *γ* = 0. When time is rescaled according to the recovery rate, the natural death rates *b*_*J*_ and *b*_*A*_ are scaled so that, when *b*_*J*_ = *b*_*A*_ = *b*, the average host lifespan 1/*b* is measured in multiples of the average duration of infection.

The host population is viable whenever *ag* − *b*_*A*_(*b*_*J*_ + *g*) > 0 and the pathogen is viable whenever the following condition holds (see *Online Resource* for derivation):

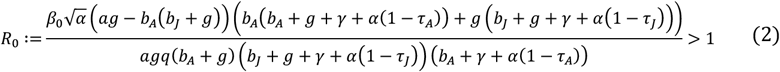

In the absence of any costs, the host will always evolve full tolerance and the pathogen will always evolve to zero virulence. We therefore assume that tolerance (juvenile or lifelong) comes at a cost to host reproduction, with the trade-off function given by:

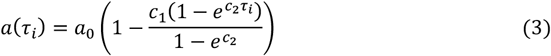

where *i* ∈ {*J, L*} depending on whether the evolving trait is juvenile or lifelong tolerance (trade-off shown in Fig. 1B). We also assume that pathogen virulence is associated with greater transmission, such that:

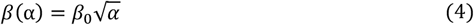

This is a commonly used trade-off function in virulence evolution models because the pathogen needs to harvest host cells in order to grow within the host and transmit (which causes damage to the host), but this process will likely have diminishing returns for the pathogen (the trade-off is shown in Fig. 1C).

### 2.2 Evolutionary dynamics

The invasion fitness of a rare pathogen mutant can be calculated using the next-generation method (see *Online Resource*) and is given by:

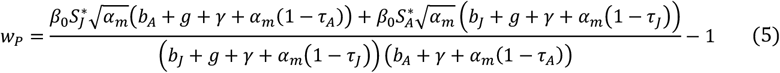

where asterisks denote the endemic equilibrium of the system.

The invasion fitness of a rare host mutant is calculated similarly (see *Online Resource*) and is given by:

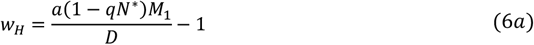

where:

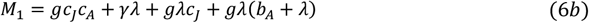

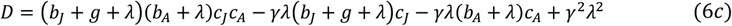

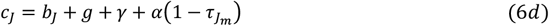

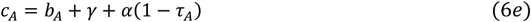

for notational convenience.

There is no closed-form analytical expression for the endemic equilibrium of system (1) or for the co-singular strategies. We therefore rely on numerical methods and simulations to determine the evolutionary endpoints of the system. To simulate the coevolution of tolerance and virulence, we first choose resident trait values and an initial population composition, both of which are arbitrary. We solve the ecological dynamics for an arbitrary, fixed time period using an ODE solver and then introduce a new, mutant sub-population of either hosts or pathogens (the individual in the current population which mutates is chosen at random, with pathogens assumed to mutate *ϕ* times faster than hosts, and the mutation is also chosen at random to increase or decrease the current trait value by a small, fixed amount). The ecological dynamics are then solved again for the same fixed time period, after which any sub-population with a sufficiently low density (again, this threshold is arbitrary) is classed as extinct and removed. A new mutant sub-population is then added as before, and these steps are repeated for many evolutionary timesteps until the long-term qualitative behaviour becomes clear.

## 3. Results

We consider the differences between our two model scenarios: lifelong and juvenile-only tolerance. Intuitively, there is a greater benefit to lifelong tolerance than to juvenile-only tolerance (because lifelong tolerance acts for longer) and so, all else being equal, lifelong tolerance will always evolve to be higher than juvenile-only tolerance. For this reason, we focus our attention on qualitative differences between the two scenarios.

A variety of different coevolutionary outcomes are possible in our model. Tolerance and virulence may both evolve to intermediate levels (at a co-CSS), may rise or fall indefinitely or to their maximum or minimum values, or may cycle indefinitely. Bistability is also possible (for a given set of parameters, we have observed anywhere from zero to three co-singular strategies, including repellers). The stability of these co-singular strategies often depends on the relative mutation rates of the host and pathogen. In particular, convergence stability may be different when the two species have equal mutation rates to when the pathogen mutates sufficiently quickly relative to the host.

### 3.1 Coevolutionary equilibria and runaway selection

In both model scenarios (when tolerance is lifelong and when tolerance is a juvenile-only trait), the host evolves to a stable level of tolerance and the pathogen to a stable level of virulence for a wide range of parameter values, with this outcome being particularly common when the host lifespan is short (Fig. S1 & Fig. S2). Both scenarios can also generate bistability (due to the presence of a repeller). In the juvenile tolerance scenario, this is most common when the pathogen baseline transmissibility is high (Fig. S1), whereas in the lifelong tolerance scenario it is most common when the pathogen baseline transmissibility is low (Fig. S2).

However, there is a significant difference between the two model scenarios when the host evolves full tolerance (*τ*_*L*_ = 1). In the lifelong tolerance scenario, this causes the pathogen to experience runaway selection for transmissibility as there is no longer an effective trade-off with virulence (virulence has no negative consequences if the host is fully tolerant throughout its lifetime). If, however, tolerance is a juvenile-only trait (and so adults are never tolerant) or if the costs of tolerance are sufficiently high (and so *τ*_*L*_ does not evolve to one) then selection for pathogen transmissibility is constrained by the negative effects of virulence. Therefore, runaway selection for increased virulence can only occur in the lifelong tolerance scenario, and only if the cost of high tolerance, *c*_1_, is sufficiently low (Fig. S2).

Tolerance (whether juvenile or lifelong) generally rises with increasing host lifespan (Fig. 2A), all else being equal. This is the result of two factors. Firstly, when hosts have very short lifespans, the infectious period is more constrained and so disease prevalence is low (and hence there is little selection for tolerance); as the host lifespan increases, disease prevalence rises which in turn increases selection for tolerance (Fig. S3A). However, if we vary the baseline pathogen transmissibility, *β*_0_, alongside lifespan in such a way that the disease prevalence is fixed, then we still see tolerance increase with lifespan (Fig. S3B). Therefore, changes in disease prevalence contribute to, but are not solely responsible for, the increase in tolerance with lifespan. Secondly, hosts with longer lifespans have more to gain from surviving an infection because they are less likely to die in any given time period than hosts with shorter lifespans. Hosts with longer lifespans therefore experience stronger selection for tolerance, no matter which model scenario is used.

**Fig 2.**
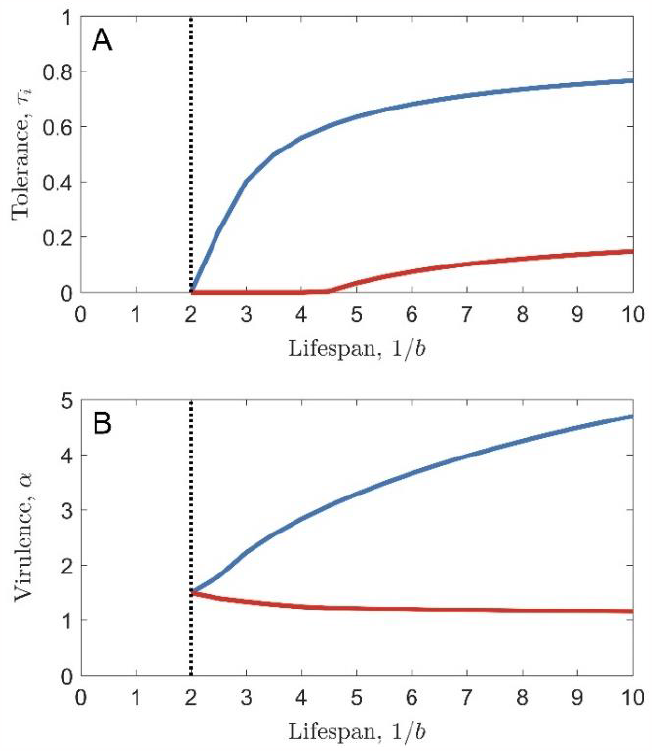
The effect of lifespan on tolerance-virulence coevolution, when tolerance is lifelong (blue curves) or limited to juveniles (red curves). The black, dashed line shows the lifespan below which the pathogen goes extinct. Parameters used are as in Table 1, with *γ* = 1 (so lifespan is measured in multiples of the average duration of infection), except for *a*_0_ = 1, *c*_2_ = 4 and *c*_1_ = 1 (full tolerance causes full sterility in the host). Results hold for all host and pathogen mutation rates (all values of *ϕ*).

The effects of lifespan on pathogen virulence, however, are dependent on whether tolerance is lifelong or restricted to juveniles. When tolerance is lifelong, virulence always increases with lifespan, whereas when only juvenile tolerance can evolve, virulence may fall with lifespan (Fig. 2B). This difference arises because of the conflicting effects of lifespan and tolerance on the evolution of virulence. If host tolerance is held constant, pathogen virulence always falls with lifespan (because there is less benefit to the pathogen to reduce its virulence if the host has a shorter lifespan and so is more likely to die soon anyway). However, if host tolerance is under selection, then longer lifespans promote higher tolerance (due to the reasons described above). Higher tolerance should therefore promote the evolution of higher virulence (because tolerance reduces the negative consequences of virulence for the pathogen by preventing the death of the host). Which of these processes dominates will determine whether virulence rises or falls with increasing lifespan. In particular, we would expect that the effect of tolerance on virulence would be stronger when lifelong tolerance evolves than when juvenile tolerance evolves, and so we would expect virulence to rise with increasing lifespan more often in the lifelong tolerance scenario than in the juvenile tolerance scenario (e.g. Fig. 2).

### 3.2 Coevolutionary cycling

As in previous models of tolerance-virulence coevolution (none of which have included age-structure, as far as we are aware), we find that coevolutionary cycling (fluctuating selection dynamics) does not occur when tolerance is constant across the host lifespan. However, when tolerance is a juvenile-only trait, coevolutionary cycling is common (Fig. 3), particularly when the host has an intermediate lifespan (Fig. 4) and when the pathogen evolves sufficiently quickly relative to the host (Fig. S4).

**Fig 3.**
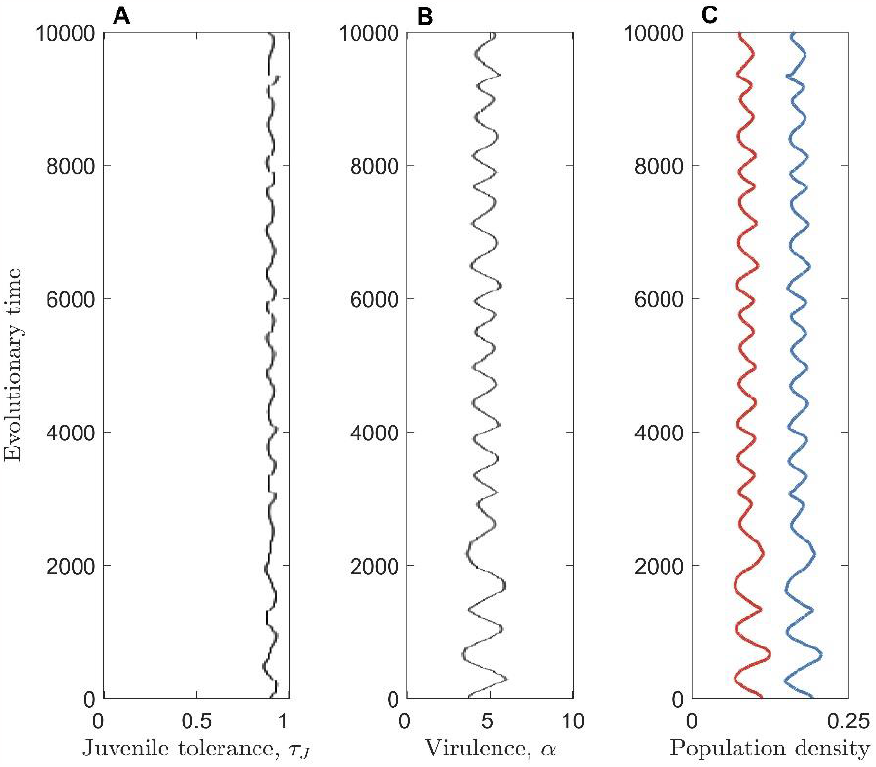
Simulation showing cycling in the host juvenile tolerance (A) and pathogen virulence (B). The total host (blue) and pathogen (red) population densities are also shown (C). Parameters used are as in Table 1, with *γ* = 0.

**Fig 4.**
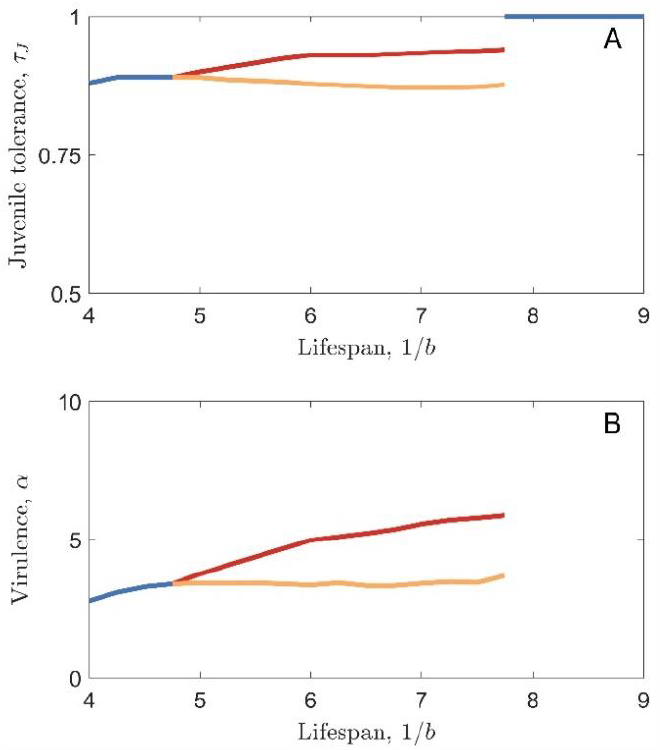
Bifurcation diagram showing the effect of lifespan on cycling. When cycling occurs, red curves show the upper limit of the cycles and orange curves show the lower limit of the cycles. Blue curves show the evolutionary endpoint in the case where no cycling occurs. Note that, when full tolerance evolves (*τ*_*J*_ = 1), virulence increases indefinitely. Parameters used are as in Table 1, with *γ* = 0, except for *c*_1_ = 0.275.

Cycling occurs in the juvenile tolerance scenario because parasitism initially selects for host tolerance to increase. As tolerance rises, virulence is no longer as detrimental to the pathogen and so selection acts to increase pathogen virulence too. High virulence causes a reduction in the density of infected hosts, because pathogens are killing their hosts (particularly the adult hosts) more quickly and so have less opportunity to transmit. A reduction in disease prevalence reduces selection for tolerance, which in turn leads to an increase in the cost of virulence for the pathogen. Virulence therefore falls, leading to an increase in the density of infected hosts, which in turn selects for increased host tolerance. The cycle then begins again. Note that cycling only occurs if higher virulence eventually causes a reduction in disease prevalence. This may seem counterintuitive, as increased virulence is accompanied by increased transmissibility. However, increased virulence also reduces the infectious period and so, unless tolerance keeps pace with virulence, disease prevalence will eventually fall, allowing cycling to occur.

Cycling occurs in the juvenile tolerance scenario and not in the lifelong tolerance scenario because increasing pathogen virulence causes a much greater reduction in the density of infected hosts in the juvenile tolerance case (because all adult hosts lack tolerance and so are strongly affected by changes in pathogen virulence). It is worth noting that the cycling we have observed is inherent to this model (not just the effect of stochasticity) and occurs via a Hopf bifurcation as lifespan varies (Fig. S5, Fig. S6). Cycling can occur both when recovery from infection is possible (*γ* = 1) and when there is no recovery (*γ* = 0), as shown in Fig. S1.

## 4. Discussion

Defences against pathogens and parasites vary with host age and yet many models assume that traits such as tolerance and resistance are consistent throughout the host’s lifetime. Here, we examine the effect of lifelong versus age-structured tolerance in a model of tolerance-virulence coevolution. We find that the life stage(s) in which tolerance acts has a significant impact on coevolutionary outcomes. In particular, coevolutionary cycling appears only to occur when tolerance can only evolve in juveniles (rather than as a lifelong trait). Previous models of non-age-structured tolerance-virulence coevolution have never observed such cycling, suggesting that age-structure may be essential for generating fluctuating selection dynamics in these models.

Age-specific tolerance also qualitatively changes the effects of host lifespan on selection for tolerance and virulence. Under all circumstances, we find that host tolerance rises with increasing lifespan, which concurs with empirical findings (Shukla et al. 2018), as well as with previous theoretical results in non-age-structured populations (Best et al. 2014). However, the effect of lifespan on the evolution of pathogen virulence depends on whether tolerance is age-restricted or lifelong, with virulence generally falling with increasing lifespan when tolerance is limited to juveniles and rising with lifespan when tolerance is lifelong.

We focussed our analysis on juvenile versus lifelong tolerance, and did not consider adult-only evolution of tolerance. There is empirical evidence that defences against disease (measured by assessing symptoms of plants exposed to pathogens and therefore reflecting the overall effect of tolerance and resistance strategies) can evolve independently at juvenile and adult stages (Bruns et al. 2022). It is therefore not unreasonable to suppose that juvenile tolerance could evolve without having a significant impact on tolerance at the adult stage. However, it may be more realistic to allow adult tolerance to evolve as well. Our model could be extended in this way to consider the coevolution of three traits: juvenile and adult tolerance and pathogen virulence. This would complicate the analysis considerably but would provide a more general picture of the effect of age-structured tolerance.

Our model has assumed a universal system of infection genetics: all pathogens can kill all hosts; pathogens with a high value of *α* are universally more virulent than those with a low value of *α*, no matter what host they are infecting; and hosts with a high value of *τ*_*i*_ are universally more tolerant than those with a low value of *τ*_*i*_, not matter what pathogen they are infected by. However, many host-pathogen systems exhibit genetic specificity, where some pathogens are better at infecting some hosts than others (Antonovics et al. 2002; Dybdahl et al. 2014). Our model could readily be extended to consider how specificity mediates selection for tolerance and virulence.

In this paper, we have considered only tolerance as a mechanism of host defence against pathogens. However, both tolerance and resistance play important roles in shaping host-pathogen interactions (Schneider and Ayres 2008). Non-age-structured, three-trait coevolution of pathogen virulence, host tolerance and host resistance has been modelled previously (Carval and Ferriere 2010), but this has never been approached in an age-structured context. Carval and Ferriere (2010) found that three-trait coevolution led to higher virulence and lower tolerance than when only one of resistance or tolerance could evolve in the host. Adding an evolvable host resistance trait to our model may have a similar quantitative effect, but the implications for our qualitative results are unclear. We have also only considered the case of mortality virulence and mortality tolerance. However, many pathogens reduce the fecundity of their hosts, either instead of or in addition to increasing mortality. Our model could be adapted to consider the coevolution of sterility virulence and sterility tolerance instead of, or in addition to, mortality virulence and mortality tolerance.

Our model incorporated two trade-offs: one between pathogen virulence and transmissibility and another between host tolerance and reproduction. Several empirical studies have found direct or indirect (via pathogen load) relationships between pathogen virulence and transmission (De Roode et al. 2008; Blanquart et al. 2016; Hawley et al. 2023), and these have inspired the virulence-transmission trade-offs used in many models of pathogen evolution, including our model. The trade-off between host tolerance and reproduction was chosen based on empirical evidence for the existence of trade-offs between disease resistance and reproduction in plants (Chaplin and Mann 1978; Simons 1979; Tian et al. 2003). However, it is possible that tolerance may come at a cost to other host life-history traits, such us maturation (*g*) or mortality (*b*). We have previously shown that the specific traits involved in trade-offs with resistance can have significant impacts on evolutionary outcomes (Buckingham et al. 2023) but whether this holds for tolerance remains to be seen.

Overall, we have shown that the life-stage(s) at which tolerance acts can have a significant impact on host-pathogen coevolutionary dynamics, in particular leading to coevolutionary cycling in tolerance and virulence when tolerance only occurs at the juvenile stage. Our findings further highlight the importance of age-structure in mediating host and pathogen evolutionary outcomes.

## Supporting information

Online Resource

## Statements and declarations

The authors declare that they have no conflict of interests.

## Data availability statement

Source code is available in the GitHub repository at: https://github.com/ecoevotheory/Buckingham_and_Ashby_2024

## Acknowledgements

Ben Ashby is supported by the Natural Environment Research Council (grant no. NE/V003909/1). This research was generously supported by a Milner Scholarship PhD grant to Lydia Buckingham from The Evolution Education Trust. We acknowledge the support of the Natural Sciences and Engineering Research Council of Canada (NSERC). Nous remercions le Conseil de recherches en sciences naturelles et en génie du Canada (CRSNG) de son soutien. The Pacific Institute on Pathogens, Pandemics and Society receives funding from the BC Ministry of Health.

