## Supplementary material for "Coevolution of age-structured tolerance and virulence": Online Resource

### The Coevolution of Age-Structured Tolerance and Virulence: Online Resource

#### Non-Dimensionalisation

We can scale the system of equations (1a) to (1d) in the main text as follows:

$$S_J, S_A, I_J, I_A, N \sim \frac{1}{q} \quad (S1a)$$

$$t \sim \frac{1}{\gamma} \quad (S1b)$$

$$a, \alpha, g, b_J, b_A \sim \gamma \quad (S1c)$$

$$\beta \sim q\gamma \quad (S1d)$$

which allows us to set $q = 1$ and $\gamma = 1$ without loss of generality.

Note that scaling by $\gamma$ is not possible in the case when there is no recovery ($\gamma = 0$). In this case, we only non-dimensionalise to set $q = 1$.

#### Invasion fitness

The ecological dynamics for a rare parasite mutant with virulence $\alpha_m$ are described by the following equations:

$$\frac{d}{dt}\begin{pmatrix} I_J \\ I_A \end{pmatrix} = \begin{pmatrix} \beta_0\sqrt{\alpha_m}S_J^* - b_J - \gamma - g - \alpha_m(1-\tau_J) & \beta_0\sqrt{\alpha_m}S_J^* \\ g + \beta_0\sqrt{\alpha_m}S_A^* & \beta_0\sqrt{\alpha_m}S_A^* - b_A - \gamma - \alpha_m(1-\tau_A) \end{pmatrix}\begin{pmatrix} I_J \\ I_A \end{pmatrix} \quad (S2)$$

The Jacobian matrix (given above) can be decomposed as:

$$J = \begin{pmatrix} \beta_0\sqrt{\alpha_m}S_J^* & \beta_0\sqrt{\alpha_m}S_J^* \\ \beta_0\sqrt{\alpha_m}S_A^* & \beta_0\sqrt{\alpha_m}S_A^* \end{pmatrix} - \begin{pmatrix} b_J + \gamma + g + \alpha_m(1-\tau_J) & 0 \\ -g & b_A + \gamma + \alpha_m(1-\tau_A) \end{pmatrix} \quad (S3)$$

Writing this decomposition as $J = F - V$, the next-generation matrix [1] is given by $N_G = FV^{-1}$ and can be written as:

$$N_G = \frac{\beta_0\sqrt{\alpha_m}}{(b_J + g + \alpha_m(1-\tau_J) + \gamma)(b_A + \alpha_m(1-\tau_A) + \gamma)}\begin{pmatrix} S_J^*(b_A + \gamma + g + \alpha_m(1-\tau_A)) & S_J^*\left(b_J + \gamma + g + \alpha_m(1-\tau_J)\right) \\ S_A^*(b_A + \gamma + g + \alpha_m(1-\tau_A)) & S_A^*\left(b_J + \gamma + g + \alpha_m(1-\tau_J)\right) \end{pmatrix} \quad (S4)$$

The invasion fitness of a rare parasite mutant is one less than the spectral radius of $N_G$. This is given by:

$$w_P = \frac{\beta_0 S_J^*\sqrt{\alpha_m}(b_A + g + \gamma + \alpha_m(1-\tau_A)) + \beta_0 S_A^*\sqrt{\alpha_m}\left(b_J + g + \gamma + \alpha_m(1-\tau_J)\right)}{\left(b_J + g + \gamma + \alpha_m(1-\tau_J)\right)(b_A + \gamma + \alpha_m(1-\tau_A))} - 1 \quad (S5)$$

The ecological dynamics of a rare host mutant with juvenile tolerance $\tau_{J_m}$ are described by the following equations (in the case where lifelong tolerance is evolving, simply replace $\tau_{J_m}$ and $\tau_A$ by the mutant lifelong tolerance, $\tau_{L_m}$):

$$\frac{d}{dt}\begin{pmatrix} S_J \\ S_A \\ I_J \\ I_A \end{pmatrix} = \begin{pmatrix} -b_J - g - \lambda & a(1-qN^*) & \gamma & a(1-qN^*) \\ g & -b_A - \lambda & 0 & \gamma \\ \lambda & 0 & -b_J - g - \alpha(1-\tau_{J_m}) - \gamma & 0 \\ 0 & \lambda & g & -b_A - \alpha(1-\tau_A) - \gamma \end{pmatrix}\begin{pmatrix} S_J \\ S_A \\ I_J \\ I_A \end{pmatrix} \quad (S6)$$

This Jacobian matrix can be decomposed as:

$$J = \begin{pmatrix} 0 & a(1-qN^*) & 0 & a(1-qN^*) \\ 0 & 0 & 0 & 0 \\ 0 & 0 & 0 & 0 \\ 0 & 0 & 0 & 0 \end{pmatrix} - \begin{pmatrix} b_J + g + \lambda & 0 & -\gamma & 0 \\ -g & b_A + \lambda & 0 & -\gamma \\ -\lambda & 0 & b_J + g + \alpha(1-\tau_{J_m}) + \gamma & 0 \\ 0 & -\lambda & -g & b_A + \alpha(1-\tau_A) + \gamma \end{pmatrix} \quad (S7)$$

We can then calculate the next-generation matrix and take one less than its spectral radius to give the invasion fitness of a rare host mutant:

$$w_H = \frac{a(1-qN^*)M_1}{D} - 1 \quad (S8a)$$

where:

$$M_1 = gc_Jc_A + \gamma\lambda + g\lambda c_J + g\lambda(b_A + \lambda) \quad (S8b)$$

$$D = (b_J + g + \lambda)(b_A + \lambda)c_Jc_A - \gamma\lambda(b_J + g + \lambda)c_J - \gamma\lambda(b_A + \lambda)c_A + \gamma^2\lambda^2 \quad (S8c)$$

$$c_J = b_J + g + \gamma + \alpha(1 - \tau_{J_m}) \quad (S8d)$$

$$c_A = b_A + \gamma + \alpha(1 - \tau_A) \quad (S8e)$$

#### DERIVATION OF $R_0$

The disease-free equilibrium of the system is given by:

$$S_J^* = \frac{b_A\left(ag - b_A(b_J + g)\right)}{agq(b_A + g)} \quad (S9a)$$

$$S_A^* = \frac{ag - b_A(b_J + g)}{aq(b_A + g)} \quad (S9b)$$

This tells us that the host population is viable whenever $ag - b_A(b_J + g) > 0$.

We can now also write the basic reproductive ratio of the parasite as:

$$R_0 = \frac{\beta_0\sqrt{\alpha}\left(ag - b_A(b_J + g)\right)\left(b_A(b_A + g + \gamma + \alpha(1-\tau_A)) + g\left(b_J + g + \gamma + \alpha(1-\tau_J)\right)\right)}{agq(b_A + g)\left(b_J + g + \gamma + \alpha(1-\tau_J)\right)(b_A + \gamma + \alpha(1-\tau_A))} \quad (S10)$$

#### **HOPF BIFURCATION**

The coevolution of host tolerance $\tau_i$ and parasite virulence $\alpha$ is described by the system:

$$\frac{d\tau_i}{dt} = \phi_H N^* \frac{\partial w_H}{\partial \tau_{i_m}}\bigg|_{\tau_{i_m}=\tau_i} \qquad (S11a)$$

$$\frac{d\alpha}{dt} = \phi_P I^* \frac{\partial w_P}{\partial \alpha_m}\bigg|_{\alpha_m=\alpha} \qquad (S11b)$$

where $\phi_H$ and $\phi_P$ are the host and parasite mutation rates respectively (and so $\phi = \phi_P/\phi_H$) and $N^*$ and $I^*$ are the endemic equilibrium host and parasite population densities.

If we let $(\tau_i^*, \alpha^*)$ represent a co-singular strategy, then we can Taylor expand the fitness gradients above to get (neglecting quadratic and higher order terms):

$$\frac{\partial w_H}{\partial \tau_{i_m}}|_{\tau_{i_m}=\tau_i} = \left(\frac{\partial^2 w_H}{\partial \tau_{i_m}^2} + \frac{\partial^2 w_H}{\partial \tau_{i_m}\partial \tau_i}\right)\bigg|_{\tau_i^*,\alpha^*} (\tau_i - \tau_i^*) + \frac{\partial^2 w_H}{\partial \tau_{i_m}\partial \alpha_m}\bigg|_{\tau_i^*,\alpha^*} (\alpha - \alpha^*) \qquad (S12a)$$

$$\frac{\partial w_P}{\partial \alpha_m}|_{\alpha_m=\alpha} = \left(\frac{\partial^2 w_P}{\partial \alpha_m^2} + \frac{\partial^2 w_P}{\partial \alpha_m\partial \alpha}\right)\bigg|_{\tau_i^*,\alpha^*} (\alpha - \alpha^*) + \frac{\partial^2 w_P}{\partial \alpha_m\partial \tau_{i_m}}\bigg|_{\tau_i^*,\alpha^*} (\tau_i - \tau_i^*) \qquad (S12b)$$

The full system can therefore be written as:

$$\frac{d}{dt}\begin{pmatrix}\tau_i - \tau_i^* \\ \alpha - \alpha^*\end{pmatrix} = \begin{pmatrix} \phi_H N^* \left(\frac{\partial^2 w_H}{\partial \tau_{i_m}^2} + \frac{\partial^2 w_H}{\partial \tau_{i_m}\partial \tau_i}\right) & \phi_H N^* \frac{\partial^2 w_H}{\partial \tau_{i_m}\partial \alpha_m} \\ \phi_P I^* \frac{\partial^2 w_P}{\partial \alpha_m \partial \tau_{i_m}} & \phi_P I^* \left(\frac{\partial^2 w_P}{\partial \alpha_m^2} + \frac{\partial^2 w_P}{\partial \alpha_m\partial \alpha}\right)\end{pmatrix}\begin{pmatrix}\tau_i - \tau_i^* \\ \alpha - \alpha^*\end{pmatrix} \qquad (S13)$$

where all derivatives are evaluated at the co-singular strategy.

A Hopf bifurcation occurs as a parameter is varied if, for some value of that parameter, the two eigenvalues of the Jacobian matrix above are purely imaginary and non-zero. In our model, a Hopf bifurcation can occur as lifespan varies (see Fig. S6).

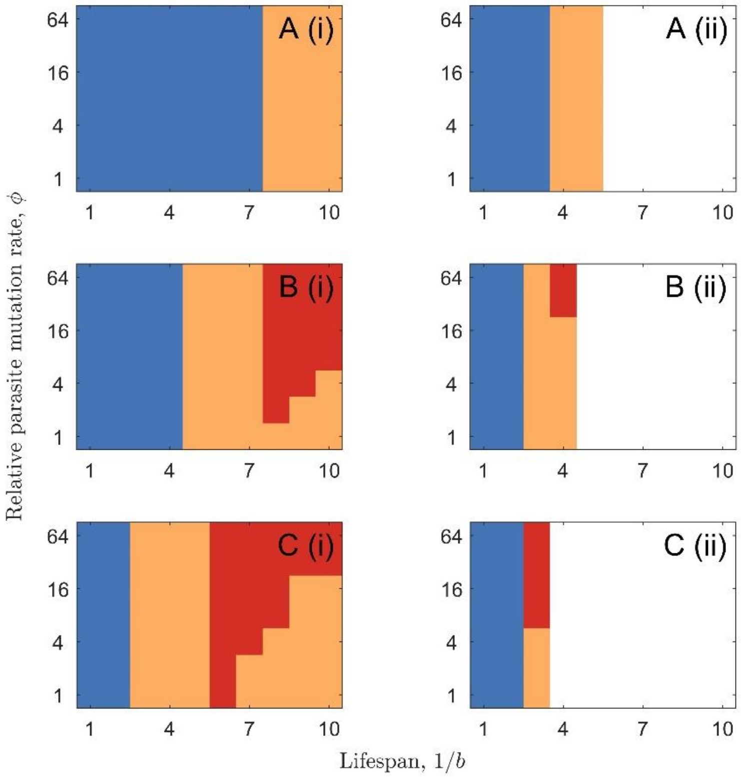


Fig. S1: The effect of lifespan and relative parasite mutation rate on the incidence of cycling and bistability in the juvenile tolerance scenario. Blue indicates a co-CSS, orange indicates bistability, red indicates cycling and white indicates that the parasite is not viable. Note that full tolerance and runaway selection for virulence never occurs. Parameters used are as in Table 1, except for $c_1 = 0.3$ and (A) $\beta_0 = 7$, (B) $\beta_0 = 10$ & (C) $\beta_0 = 20$, with (i) $\gamma = 0$ & (ii) $\gamma = 1$.

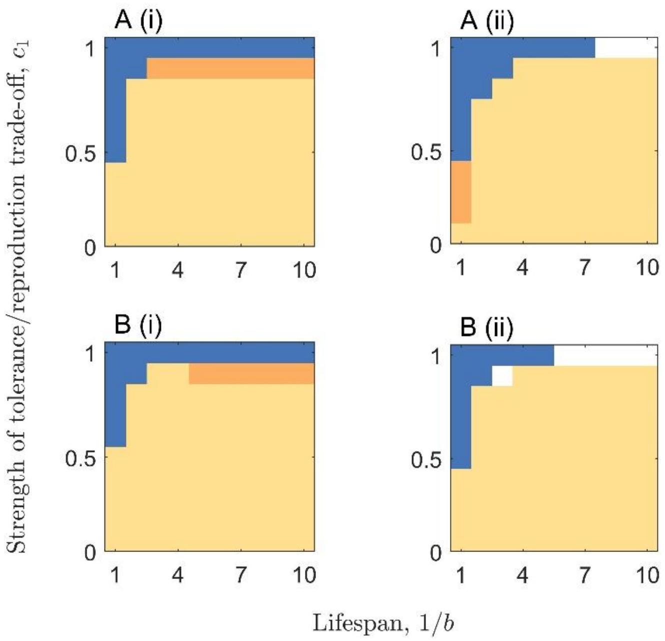


Fig. S2: The effect of lifespan and the strength of the tolerance/reproduction trade-off on the incidence of bistability and the evolution of full tolerance in the lifelong tolerance scenario. Blue indicates a co-CSS, orange indicates bistability, yellow indicates the evolution of full tolerance and runaway selection for virulence and white indicates that the parasite is not viable. Note that cycling never occurs. Parameters used are as in Table 1, except for (A) $\beta_0 = 10$ & (B) $\beta_0 = 20$, with (i) $\gamma = 0$ & (ii) $\gamma = 1$. Results hold for all host and parasite mutation rates (all values of $\phi$).

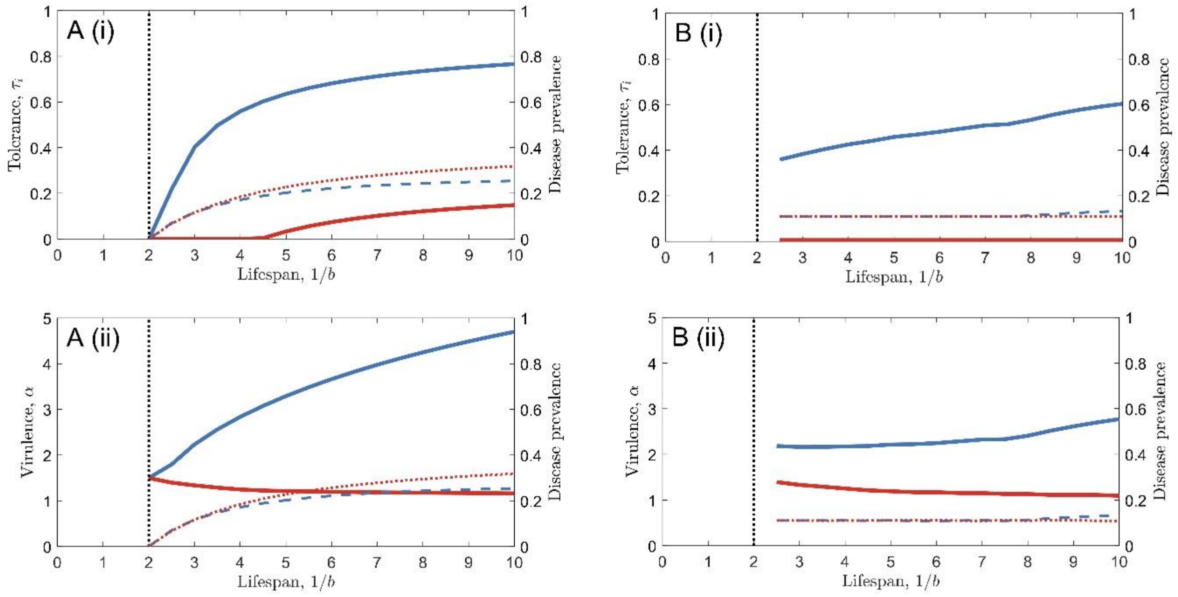


Fig. S3: The effect of lifespan on (i) tolerance and (ii) virulence in the case of evolving lifelong tolerance (solid blue curves) and evolving juvenile tolerance (solid red curves). The black, dashed line shows the lifespan below which the parasite goes extinct. The blue and red dashed lines show the disease prevalence at the co-singular strategy in the lifelong and juvenile tolerance scenarios respectively. In (A), $\beta_0$ is held fixed throughout, whereas in (B), a different value of $\beta_0$ is chosen for each value of lifespan to ensure that the disease prevalence is fixed. Parameters used are as in Table 1, with $\gamma = 1$ (so lifespan is measured in multiples of the average duration of infection), except for $a_0 = 1$, $c_2 = 4$ and $c_1 = 1$ (full tolerance causes full sterility in the host). Results hold for all host and parasite mutation rates (all values of $\phi$).

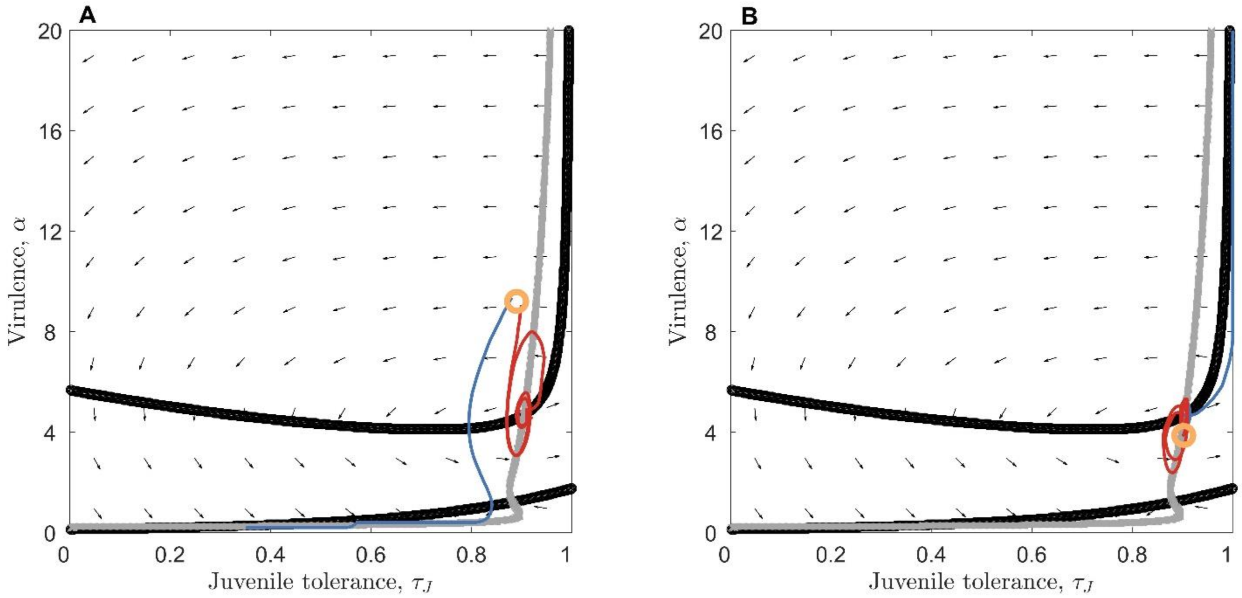


Fig S4: Phase planes showing coevolution for different relative mutation rates. The host tolerance nullcline is shown in black and the parasite virulence nullcline is shown in grey. Arrows show the directions of the fitness gradients (in the case where $\phi = 1$). Trajectories (initiated at different points in (A) and (B), as indicated by the orange circles) are shown in blue (when the host and parasite have equal mutation rates; $\phi = 1$) and in red (when the host mutates ten times faster than the host; $\phi = 10$). Parameters used are as in Table 1, with $\gamma = 0$.

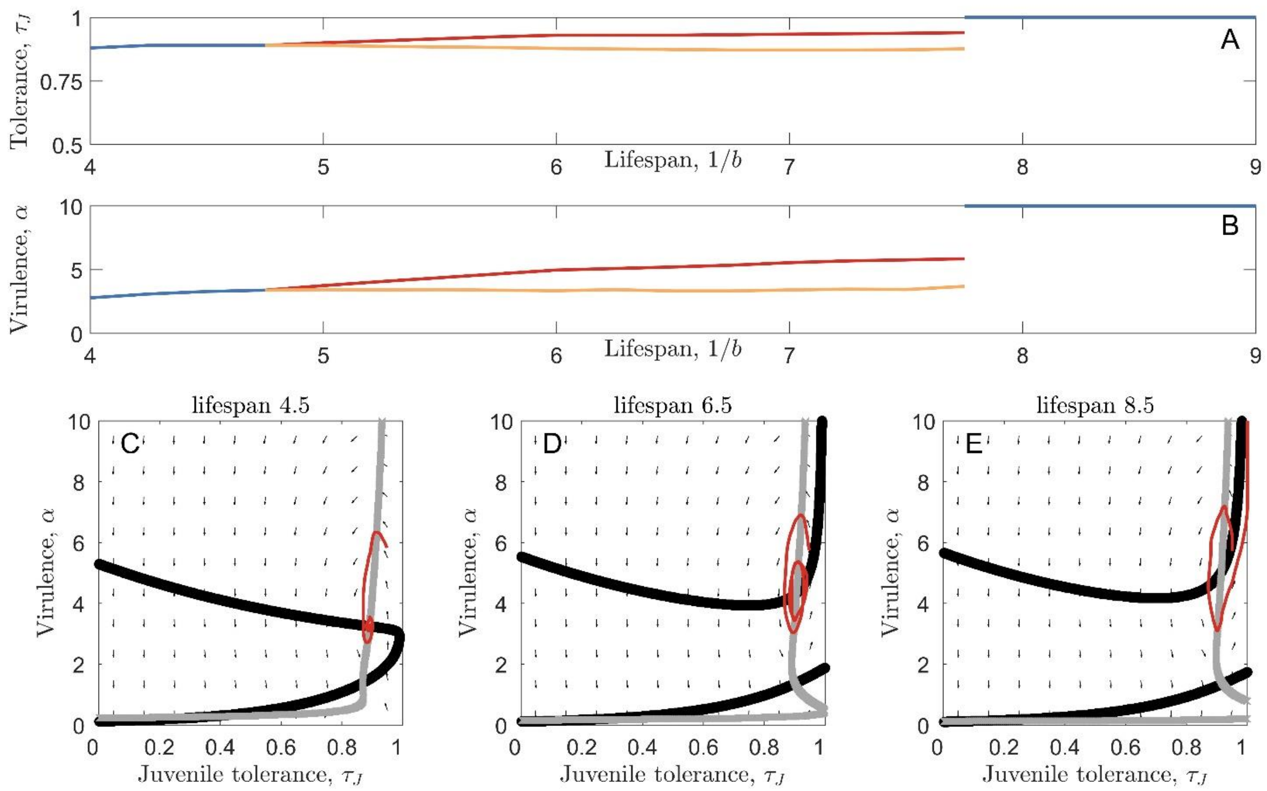


Fig. S5: Bifurcation diagrams showing the effect of lifespan on cycling in the host juvenile tolerance (A) and parasite virulence (B). Blue curves show the co-CSS when cycling does not occur; red and orange curves show the upper and lower limits of cycles when cycling does occur. Phase planes show the host and parasite nullclines (black and grey respectively), directions of fitness gradients (arrows) and a trajectory (red), in the cases of short (C), intermediate (D) and long (E) lifespans. Parameters used are as in Table 1, with $\gamma = 0$, except for $c_1 = 0.275$. Lifespans are $1/b = 4.5$ in panel (C), $1/b = 6.5$ in panel (D) and $1/b = 8.5$ in panel (E). All trajectories are initiated at $\tau_J = 0.95$ and $\alpha = 5.5$. Arrows and trajectories all reflect a relative parasite mutation rate of $\phi = 20$ (the parasite mutates 20 times faster than the host).

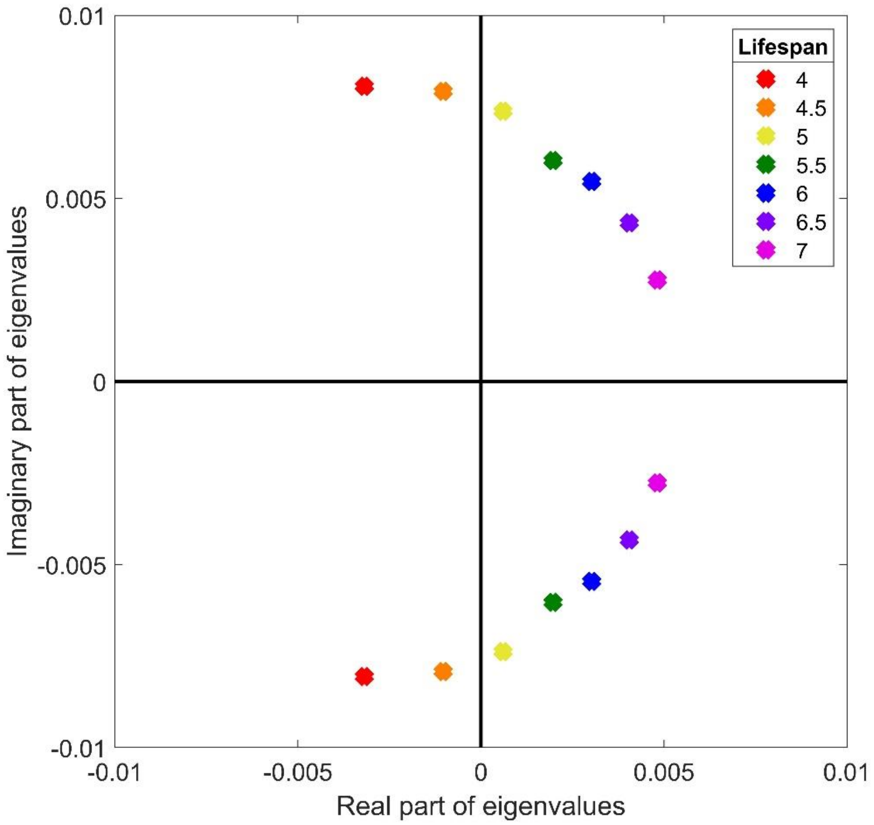


Fig. S6: Argand diagram showing the eigenvalues of the Jacobian matrix of the full coevolutionary system as lifespan varies. A Hopf bifurcation can be seen as lifespan increases (the eigenvalues cross the imaginary axis away from the origin). Lifespan ($1/b$) increases from 4 to 7 as eigenvalues move left-to-right across the plane. Parameters used are as in Table 1, with $\gamma = 0$, except for $c_1 = 0.275$.
